# Diurnal variations in the motility of algal populations

**DOI:** 10.1101/854844

**Authors:** D. Jin, J Kotar, E. Silvester, K. C. Leptos, O. A. Croze

## Abstract

The motility of microalgae has been studied extensively, particularly in model microorganisms such as *Chlamydomonas reinhardtii*. For this and other microalgal species, diurnal cycles are well-known to control the metabolism, growth and cell division. Diurnal variations, however, have been largely neglected in quantitative studies of motility. Here, we demonstrate using tracking microscopy how the motility statistics of *C. reinhardtii* are modulated by diurnal cycles. We discovered that the mean swimming speed is greater during the dark period of a diurnal cycle. From this measurement, using a hydrodynamic power balance, we conjecture that this is a result of the mean flagellar beat frequency being modulated by the flagellar ATP. Our measurements also quantify the diurnal variations of the orientational and gravitactic transport of *C. reinhardtii*. We discuss the implications of our frequency results in the context of cellular bioenergetics. Further, we explore the population-level consequences of diurnal variations of motility statistics by evaluating a prediction for how the gravitactic steady state changes with time during a diurnal cycle.

**SIGNIFICANCE:** We report tracking microscopy measurements which demonstrate that the mean swimming speed of *C. reinhardtii* is significantly greater during the dark period of a diurnal cycle. Using hydrodynamic (low Reynolds number) power balance, we also inferred the mean flagellar beat frequency from the swimming speed, hypothesising that the observed variations in this frequency correlate with the diurnal regulation of flagellar ATP. Diurnal variations of the orientational and gravitactic transport of *C. reinhardtii* were also quantified and used in a continuum model to predict that, at the population scale, the steady state vertical distribution of *C. reinhardtii* is broader during the dark period. Our findings could have significant implications for microalgal biotechnologies, e.g. microalgal harvesting, and plankton migration in the ocean.

## INTRODUCTION

Many microorganisms experience changes in light and darkness corresponding to the regular alternation of day and night, and have evolved circadian clocks to adapt to these environmental rhythms (1). The effect of these diurnal cycles is particularly important for photosynthetic microorganisms, such as the model unicellular microalga *Chlamydomonas reinhardtii*, for which light is only available for growth during the day. For *C. reinhardtii*, diurnal cycles have been shown to set endogenous circadian clocks (2), allowing for the optimal distribution of metabolic processes (3), growth and division(4) between light and dark phases.

*C. reinhardtii* swims by beating a pair of flagella. The study of its motility, from the molecular details of flagellar motion through to the collective behaviour of whole populations(5–7), continues to be a very active area of study. While it is known that the circadian clock of *C. reinhardtii* regulates phototaxis (8) and chemotaxis in zoospores (9), the swimming motility of *C. reinhardtii* has not been comprehensively quantified as a function of diurnal cycles, as we do in this study.

In previous studies measuring the swimming speed of *C. reinhardtii*(10–16), cells were usually sampled at a single time point in the cell cycle under growth conditions (temperature, growth medium, and illumination) not corresponding to those necessary for obtaining diurnally synchronised cultures. Similar considerations apply to quantitative measurements of motility in other microalgal species. For example, swimming speed, rotational diffusivity and gravitatic reorientation time (the time it takes a cell to gravitactically reorient to the vertical) have recently been measured for the marine species *Dunaliella salina*(17) and *Heterosigma akashiwo*(18, 19). However, like in the studies of *C. reinhardtii*, these microalgae were not synchronised to diurnal cycles.

*C. reinhardtii* flagella are resorbed during mitosis (20), so it is expected that the number of motile cells will change when cells divide. However, it is not a priori clear how other statistical measures of motility, such as mean swimming speed, rotational diffusivity and gravitatctic reorientation time, will change diurnally. In the current study, we use tracking microscopy, augumented by individual-based simulations, to obtain these motility statistics as a function of time during the cell cycle of *C. reinhardtii* cells synchronised to light-dark cycles.

## MATERIALS AND METHODS

### Culturing conditions and synchronisation of cell division

*C. reinhardtii* CC-125 wt cells were transferred from slant cultures to 130 ml Tris-minimal (TM) medium and grown axenically in 250 ml conical flasks in a shaking incubator (Infors Minitron). Cultures were constantly shaken at 100 rpm and bubbled with air at constant temperature 25 ^o^C. Cultures were exposed to a 14:10 light-dark cycle with 300 µmol/(m^2^ .s) photosynthetically active radiation. At the beginning of each light period, the culture was diluted to 5 × 10^5^ cells/ml with TM medium to keep cultures in exponential growth phase (21). Experiments were carried out after cells had reproduced for more than five generations. Experimental data was collected from nine independently innoculated and incubated flask cultures.

### Sampling, cell enumeration and imaging

At various times during the 24-hour light:dark cycle, an aliquot sample was extracted from a culture flask and motility, counts and density measurements were performed in a sequence I-V, as below, lasting at most 1.5 hours. Experiments were repeated with 9 independent flask cultures, each sampled at random time points to give a total of 31 data sets. *I. Cell counts.* 0.5 µl suspension was diluted in 10 ml of ISOTON® buffer solution and measured with a Beckman Z2 Coulter counter. Measurements were repeated three times for each sample. Each cell number density and volume measurement corresponded to more than 2500 particles. *II. Microscopy of swimmers.* Cells were imaged using a vertical-stage microscope via a 4× objective (Olympus UPlanFL N, 0.13 N.A.) with bright field deep red illumination (peaking at 660 nm to avoid eliciting a phototactic response (12, 22)), see Figure 2 a. Suspensions of cells were transferred using a micropipette to a 0.4 × 8 × 50 mm rectangular capillary, which was then loaded onto the microscope stage. The suspension was allowed to rest for 10s so fluid motion could dissipate and the distribution of the cell orientations could equilibrate, as confirmed by tracking analysis. Movies of swimming cells, each 20 s long, were captured at 50 or 162 frames per second with a CMOS camera (Grasshopper3 GS3-U3-23S6M), with a spatial resolution of 1.5 µm/pixel for the magnification employed. We recorded 10 individual videos at a cell number densities approximately 5 × 10^5^ cells/ml during the light period, and 5 individual videos at densities of 1 × 10^6^ cells/ml and above in the post-division dark period. *III. Recording non-swimmers.* The capillary was transferred to a horizontal-stage microscope (Olympus IX73). For times ≳ 2 minutes, non-swimmers (settling speed ≈ 5 µm/s) settled to the bottom of the capillary (depth of 400 µm). Focusing on bottom of the capillary, videos, containing static and swimming cells, were recorded at 1 frame per second at five different locations. *IV. Recording cell sedimentation.* See supporting information. *V. Cell shape and condition.* A few images of the immobilised cells were taken with the horizontal stage microscope using a 60× objective (Olympus LUCPlanFL N, 0.70 N.A.) to examine the shape and the physiological state of the cells.

### Quantifying motile fraction, swimming speed and shape parameters

#### Swimmer fraction

A median filter was applied to videos containing nonswimmers; this removes swimming cells. Static nonswimming cells cells were then counted using the particle identification function in the tracking algorithm (23). The number density of non-swimmers was calculated in the volume defined by the capillary depth and the area of the field of view. Finally, the swimmer fraction is evaluated as the ratio between the nonswimmer density and the total number density obtained from Coulter counter measurements. *Mean swimming speed.* A background image obtained by median filtering the time sequence of the swimming cells was subtracted from image sequences to remove static cells. Cell tracking was performed with a MATLAB implementation of the Crocker and Grier algorithm (23). From each set of videos, between 1550 to 54000 trajectories were acquired, with the higher end corresponding to the post-mitotic period. *Chlamydomonas* is known to swim helically, turning counter-clockwise at a frequency of 2 Hz (24), which manifests itself as small oscillations in the 2D projected trajectories. These oscillations were removed using a median filter at 0.25 s intervals. Consequently, 0.25 s segments were truncated from the two ends of each trajectory, (Figure 2 b). Trajectories tracked originate from a volume 200 µm in depth, of the same order of magnitude as the average trajectory length, and therefore the trajectories are 2D projections of the 3D trajectories (25). The true swimming speed is evaluated assuming that swimming speeds have a sufficiently narrow distribution, in which case, 〈*v*_s_〉^2^ ≈ 〈*v*_s_^2^〉 = 〈*v*_x_^2^〉 + 〈*v*_z_^2^〉. The assumption is reasonable for *C. reinhardtii* CC-125: for this strain, Fujita et al. (2014) observed that the variation of swimming speed is approximately 23% of the mean, and is mainly a result of cell to cell variation. The decorrelation time of the swimming speed in either direction was estimated to be 6 s (see Supporting Information), therefore, *v*_x_ and *v*_z_ were evaluated at 6 s intervals along the trajectories to ensure the mean evaluated is unbiased. Each data set consists of 5 to 10 repeated recordings, and the error is evaluated as the standard error of mean in the data set. When cells are under going mass division, they become non-motile and appear as settling cells in videos. In analysis (see below) nonmotile cells were identified and removed. *Assessing cell shape.* The major and minor axes were measured from the high resolution images manually in MATLAB. *Zeitgeber time assignment.* After analysis, the final data sets were grouped according to the time intervals in which they were acquired and represented and assigned an average time and parameter values. Sampling during the light period caused growth to be interrupted as cells were transferred to microscopy under red light. Data sets acquired under these conditions were assigned the time corresponding to sampling from the flask culture to represent the hours of light exposure more accurately. On the other hand, data sets acquired during the dark period were assigned the time of video recording.

### Gravitactic parameters, rotational diffusivity and mass density

Assuming steady-state gravitactic swimming statistics, the bias parameter *λ* can be evaluated from 〈*v*_z_〉/*v*_s_ = coth(*λ*) − 1/*λ*, where 〈*v*_z_〉 is the ensemble average of the vertical component of the swimming velocities, obtained from tracking as above, and *v*_s_ is the swimming speed (26). The rotational diffusivity *D*_r_ was inferred from the direction correlation function of the tracked data. We carried out 3D individual-based simulations, which allowed to properly deduce the value of *D*_r_ from the 2D projections available in the tracked data (see Supporting Material). The gravitactic reorientation *B* was inferred indirectly from the definition of *λ* = 1/(2*BD*_r_). The mass density of cells was deduced from the settling speed of UV immobilised *C. reinhardtii* cells (see Supporting Material).

### Theoretical prediction of vertical gravitactic distribution

In the absence of background flow, the concentration *n*(**r**, *t*) of a population of gravitactic algae obeys(26)

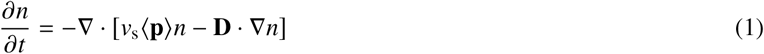

where 〈**p**〉 = (0, 0, *K*_1_) is the mean cell orientation, **D** = diag(*K*_1_, *K*_1_, *K*_2_) is the anisotropic diffusivity tensor, and *K*_1_ (*λ*) = coth (*λ*) − 1/*λ* and *K*_2_ (*λ*) = 1 − coth (*λ*) ^2^ + 1/*λ*^2^ are functions of the bias parameter *λ*, see (26, 27). As previously discussed in the paper, *v*_s_ is the cell mean swimming speed and *D*_r_ the rotational diffusivity. We seek the steady state (*∂n/∂t* = 0), so that equation (1) becomes

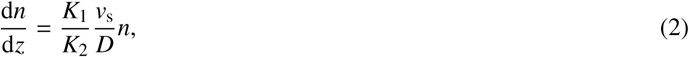

which, with a background concentration *n*_*b*_ and a free surface at *z* = *h* and requiring 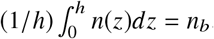, is solved by

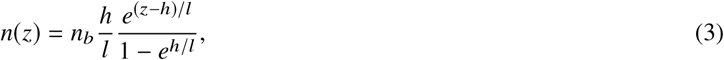

where we have defined the characteristic length scale 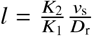.

## RESULTS

### Growth and division of synchronised swimmers

Cells of *Chlamydomonas reinhardtii* CC125, a model wild-type strain for flagellated motility (15, 28, 29), were entrained to a 14:10 light-dark cycle, as detailed in Materials and Methods. Images taken at different time points through the cell cycle, indicated as Zeitgeber Times (ZT) in hours following the onset of illumination, confirm qualitatively the synchronization of growth and reproduction of cells to the light-dark cycle (Figure 1 a-h). These were further quantified by measurements of cell size and number (see Materials and Methods). During the light period, cells grow photosynthetically, increasing in size (Figure 1 j) and becoming more spherical in shape (Figure S2 b in the Supporting Material). The cell number density in this period is stationary (Figure 1 i). Subsequently, 3-4 hours into the dark period, cells undergo mitotic division and rapidly proliferate (Figure 1 a). This is indicated by a sharp reduction of average cell volume between ZT14 to ZT15 (Figure 1 j). In the late dark period, while the cell number density approaches a steady level (Figure 1 i), cell volume continues to decrease. These observations reflect the canonical behaviour of synchronised microalgal cells. Next, we consider how motility changes through the diurnal cycle.

### Unexpected diurnal changes in swimming speed

We used tracking microscopy to characterise the diurnal variations in swimming motility of *C. reinhardtii* (see Materials and Methods). During the light period, the swimmer fraction, shown in Figure 2 c, is constant and high (≈96%). Its value then decreases at the onset of the dark period, during which the fraction of swimmers reaches a minimum at around ZT15. Subsequently, the swimmer fraction grows back to its light period value at ZT22. This pattern reflects the well-known cessation of motility of mother cells due to flagellar resorption preceding mitosis, and followed by hatching of newly-flagellated motile daughter cells. The average post-mitosis cell number density, *n* = 2.61 × 10^6^, derives from an average of 2.28 divisions of 5 × 10^5^ mother cells. Assuming 30 minutes for flagella resorption (30) and 30 minutes for each division cycle (4), the total division time is estimated to take approximately 90-150 minutes. This estimate is in good agreement with the temporal trends of the cell number density and volume discussed above, and is also consistent with the description of synchronized *Chlamydomonas* mitosis in existing studies (31). Swimming fraction results thus reflect the expected cellular dynamics of synchronised cells.

Measurements of mean swimming speed, however, reveal an unexpected and surprising trend: the speed falls from 140 − 160 µm/s during the dark period to 80 µm/s at the onset of the light period (Figure 2 d). While the order of magnitude of the swimming speed is comparable with that of previous studies (10, 14, 15), the intriguing diurnal change we have here observed has not, to the best of our knowledge, previously been reported.

**Fig. 1.**
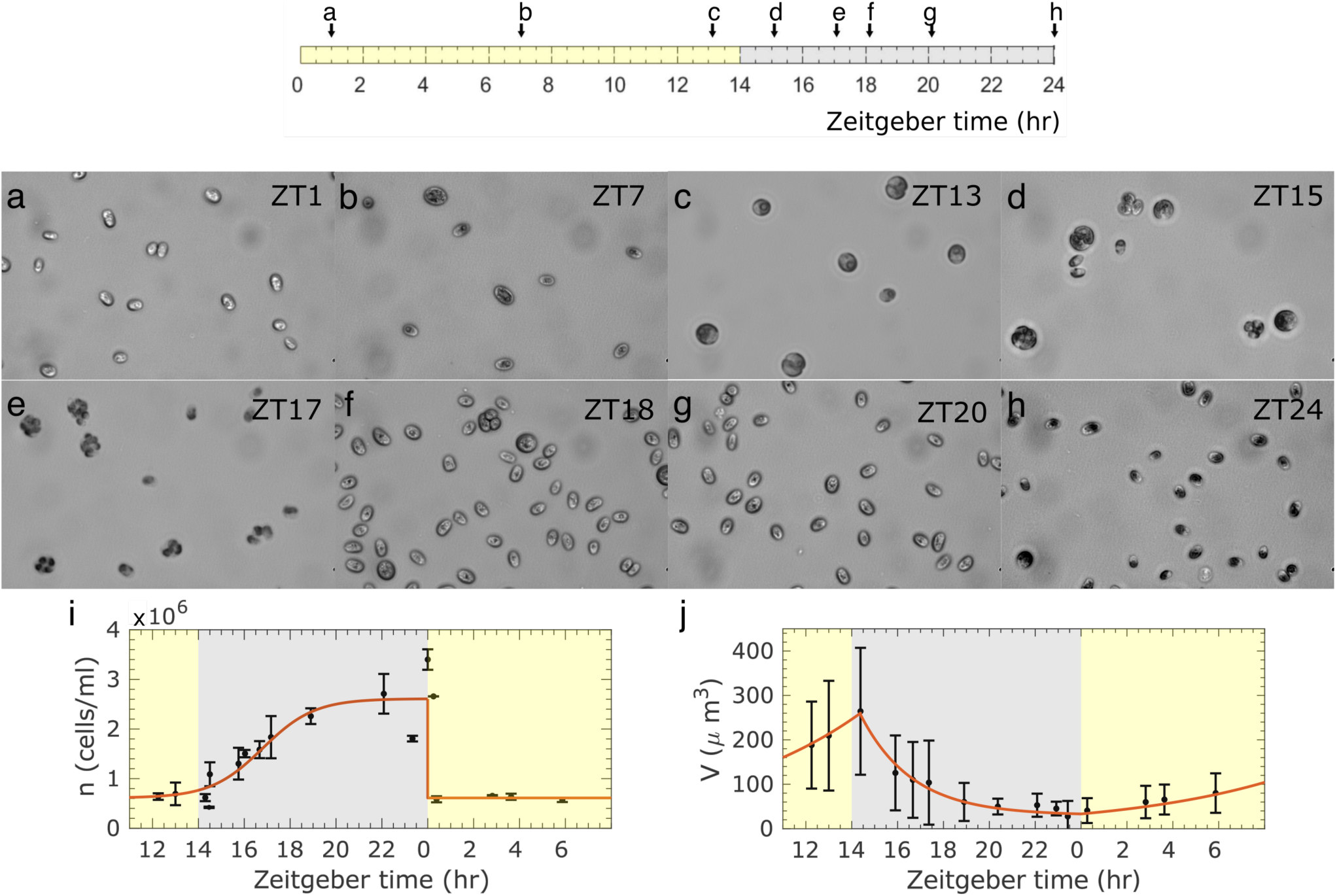
a-h) Images of *C. reinhardtii* cells at different time points in a 14:10 light-dark cycle. Note that variation in cell opacity is due to different optical contrast. i) Cell number density fitted to a logistic function *n* = *n*_*max*_/(1 + exp(−*k*(*t*−*t*_*c*_))) + *n*_0_, where *n*_*max*_ = 2 × 10^6^ cells/ml, *k* = 0.885 hr^−1^, *t*_*c*_ = 16.8 hr, *n*_0_ = 6.11 × 10^5^ cells/ml. Error bars represent standard error of mean. j) Cell volume fitted to exponential growth and decay trends, *V* = *V*_1_ exp(*k*(*t*−*t*_*c*_)) + *V*_2_, respectively with *V*_1_ = 33.2 µm^3^, *k* = 0.143 hr^−1^, *t*_*c*_ = 0, *V*_2_ = 0, and *V*_1_ = 227 µm^3^, *k* =−0.445 hr^−1^, *t*_*c*_ = 14.4 hr, *V*_2_ = 30.0 µm^3^. Error bars represent standard deviations of the volume distribution.

**Fig. 2.**
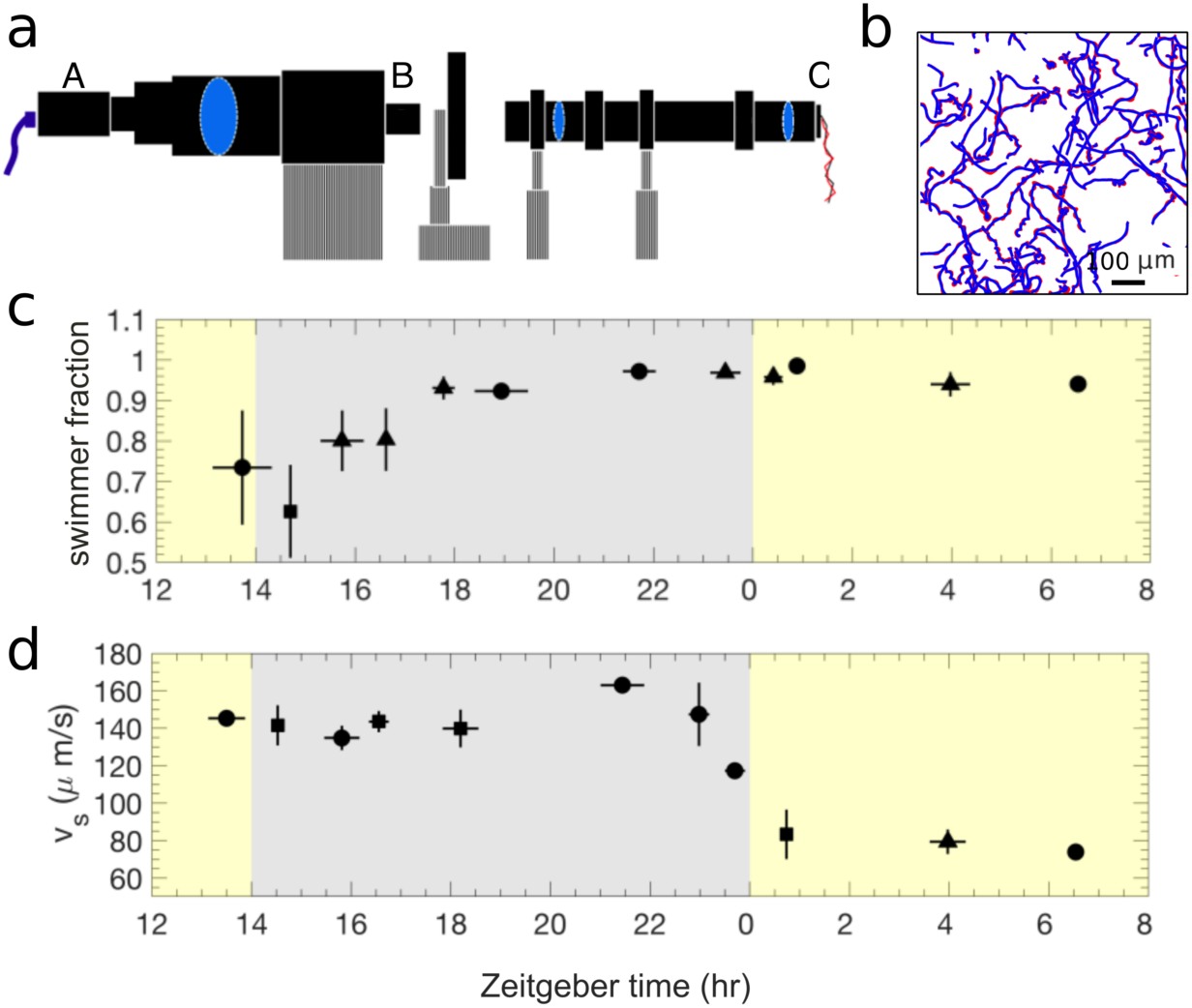
a) An illustration of the vertical-stage microscope with Köhler illumination (side view). A: camera. B: 4× objective lens. C: LED illumination. b) Original trajectories (red) were reduced to the large-scale motility dynamics by a median filter which removes the helical swimming oscillations (blue). c) Fraction of the cell population that are swimmers. d) Swimming speed. •, ▴, and ▪ symbols respectively represents a sample size of 2, 3, 4.

### Flagellar beat frequency and power

By a simple hydrodynamic argument, we can connect the diurnal variation of measured mean speed with the associated variation in mean flagellar beat frequency, which, as discussed below, can be connected to cellular bioenergetics. The beating flagella of *C. reinhardtii* transmit power to the surrounding fluid and translate the cell against the resistive drag force exerted by the fluid on the cell body. Because this swimming motion occurs at low Reynolds number (Re= *v*_s_*a*/*v* ~ 10^−4^, with values of *v*_s_ and *a* from Figures 2 and S2, respectively, and *v* ≈ 10^2^ µm/s^2^ is the dynamic viscosity for water at 25°C) we can write the propulsive power balance as:

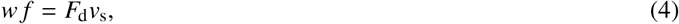

where w is the work per beat performed by flagella, *f* is the flagellar beat frequency, *v*_s_ is the swimming speed and *F*_d_ is the drag force. All quantities are averages over the beat cycle. The drag force is given by Stokes law

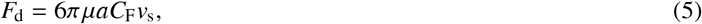

where *µ* is the kinematic viscosity of the growth medium, and the drag coefficient is given by 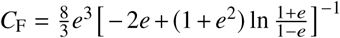 where 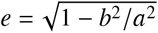 is the mean eccentricity of cell, assumed to have the shape of a spheroid with semi-major and semi-minor axes *a* and *b*, respectively. Substituting the expression for *F*_d_ from (5) into equation (4), we then obtain:

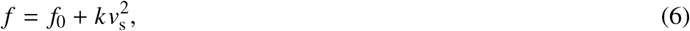

where we have defined *k* = 6*πµaC*_F_/w and *f*_0_, an experimental threshold frequency defined below. Fujita et al.(15) have measured swimming speed as a function of flagellar beat frequency of *C. reinhardtii* including CC-125, the wild-type strain used in this study, and four outer dynein arm mutants *oda*1, *oda*11, *suppf* 1, and *suppf* 2. These mutants have a breastroke motion similar to the wild type, but beat at different (lower) frequencies (15). We introduce here a threshold frequency *f*_0_ to account for the fact that, at low beating frequencies, the translational displacement of mutants falls below resolution of the tracking method used to measure swimming(15). Fitting the Fujita et al.(15) data using equation (6) gives *f*_0_ = 29 Hz and *k* = 9.6 × 10^−4^s/µm^2^.

The diurnal variation of average beat frequency thus inferred from our speed data is shown in Figure 3 a. The order of magnitude values of the beat frequencies agree with previously reported values (10, 15, 29), however we believe that the diurnal variation in the frequency is reported here for the first time. Further, using (5) and (4), we can also obtain the variation in mean drag force and power, respectively. These are shown in Figure 3 a. The power is of the same order of magnitude as the mean power inferred by Guasto *et al.* from analysis of the flow field around swimming *C. reinhardtii*(32). As other studies, however, this study did not investigate diurnally synchronised cells.

**Fig. 3.**
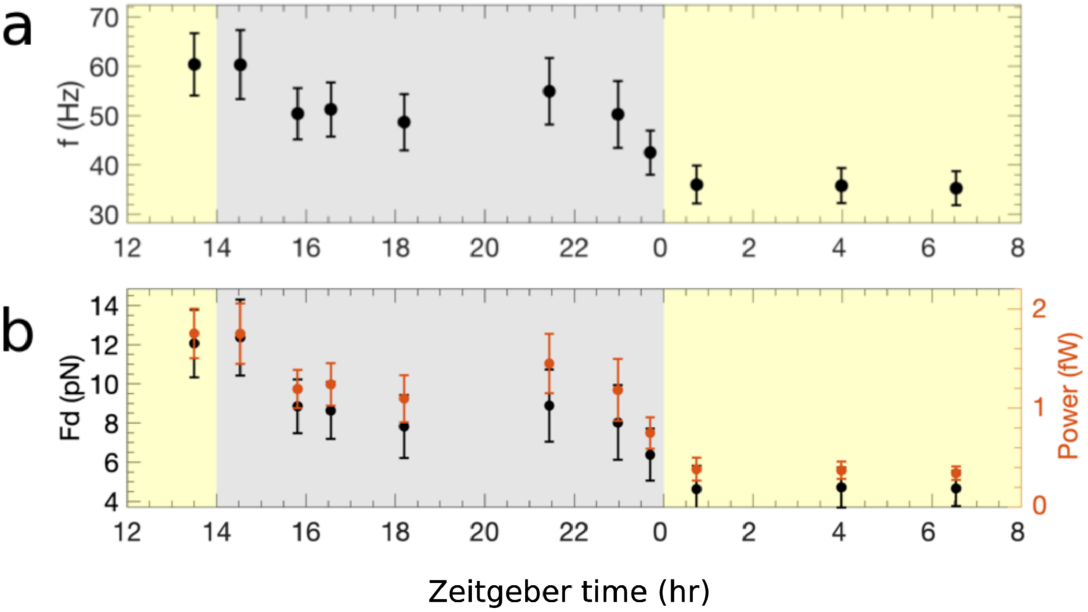
Diurnal variation of a) Average beating frequency and b) average drag force and power.

### Gravitactic parameters, rotational diffusivity and mass density

Aside from mean speed and flagellar frequency, we measured other motility and hydrodynamic parameters important for the transport and dispersion of suspensions. These measurements are important to allow the parameterisation of mathematical models of cell suspensions, as has been done for other species, but ignoring diurnal variations (17). The parameters measured include the gravitactic reorientation time *B*, and rotational diffusivity *D*_r_, and the bias parameter *λ* = 1/(2*BD*_r_), which quantifies the ratio of rotational diffusive and gravitational reorientation times. The diurnal variation of these parameters is plotted in Figure S3 c,d and a, respectively. The data for these parameters is more noisy than for the swimming speed, and it is harder to identify trends. The bias parameter *λ* is approximately constant in value during the light period and into the dark period, except for an apparent dip in value prior to the start of the light period. The reorientation time *B* appears to grow during the dark phase, falling at the start of the light phase, and growing again during the day. The rotational diffusivity *D*_r_ also grows in the dark, but appears to then fall during the day. The largest values of *B* we have measured are four times larger than what has been previously reported(10) for *C. reinhardtii*. The values for *D*_r_ are larger but of the same order of magnitude as reported in previous studies (14, 33). The bias parameter *λ* has not been previously measured before for *C. reinhardtii*, and, across the diurnal variations, is measured to be smaller than that quantified for other gravitactic species, e.g. *λ* = 0.21 for *D. salina*(17) and *λ* = 2.2 for *Chlamydomonas augustae*(25).

Tracking settling heat-immobilised cells, we were also able to quantify the mean mass density ⇢ of cells, an important parameter for the analysis of suspension bioconvective instabilities (26) and for coupling the gyrotactic dynamics of cells to flow, e.g. in pipe flows (17). As shown in Figure S4, the density appears 10% larger than the surrounding fluid, with a slight reduction in its value during the light period.

## DISCUSSION

We have characterised the motility of *C. reinhardtii* through light-dark cycles using tracking microscopy. Our results reveal a previously unobserved diurnal variation in the mean swimming speed of this model species. Further, using a hydrodynamic power balance, we were able to infer from the speed the corresponding diurnal variation in mean beat frequency. A study of demembraned *Lytechinus pictus* sea urchin sperm flagella has demonstrated ATP concentration correlates positively with flagellar beat frequency (34). We can reasonably hypothesise that a similar correlation will apply qualitatively for *C. reinhardtii*. If this is the case, then changes in flagellar frequency shown in Figure 3 a reflect changes in flagellar ATP. In particular, our data would suggest that during the dark period the amount of ATP available to flagella drops in value to a lower level that is maintained during the day; flagellar ATP then surges at the onset of division. To test if the flagellar frequency does indeed chart the variation of flagellar ATP, it would be very interesting to carry out experiments similar to (34) assaying the diurnal variation of ATP in the flagella of *C. reinhardtii*.

Our results also quantify the power exerted by the cell on the fluid when swimming, shown in Figure 3 b. This power, like the frequency, broadly decreases during the dark phase to a lower light-phase value. Across the diurnal variation, its value, ∼ 1fW, is an order of magnitude smaller than the minimum mechanistic power required for flagellar motility, estimated by Raven from the reaction rate of dynein molecules along the flagella to be 22 fW for freshwater microalgae with the same broad characteristics as *C. reinhardtii*(35). This power, in turn, was estimated to be half of the minimum cell maintenance power(35). These estimates demonstrate that swimming is energetically affordable to photosynthetic microalgae, as opposed to other microorganisms, such as bacteria, for which it represents a much more prohibitive metabolic cost (26, 36). However, order of magnitude estimates are unable to shine light on the variations in cellular bioenergetics that underly the diurnal variations in swimming we have observed.

The flagellar beat is powered by ATP in the flagellar compartment (37), which has either arrived there by diffusion from the cytosol or has been generated *in situ* by reactions with the metabolite 3-phosphoglycerate (3PG). During the photosynthetic light period, ATP and 3PG molecules are produced by the central carbon metabolism in reactions compartmentalised in the chloroplast, mitochondria, or the cytosol(38). Synthesis of energy-storing metabolites (starch, lipids), nutrient uptake, cell maintenance and growth, all consume ATP during the light period. Flagellar motion also consumes, we can assume, a small fraction of the ATP budget. We can interpret our results to suggest that ATP available to flagella surges at the onset of division. This is associated with the transition from photosynthetic growth to mitotic division powered by respiration. During the dark period, in the absence of photosynthesis, only mitochondria and the cytosol, by degrading storage products using respiration and fermentation (3), are active in producing ATP to power cell division and flagellar beating. The balance of these intracellular processes will determine how much ATP is available to flagella. Plausibly, as the total pool of ATP diminshes with the degradation of storage products, there will be less ATP available for flagella, qualitatively accounting for the decline towards the end of the dark period we infer from our results. It will be interesting in future experiments and modelling to quantify the intracellular energy budget and connect it with swimming measurements like those we have carried out.

**Fig. 4.**
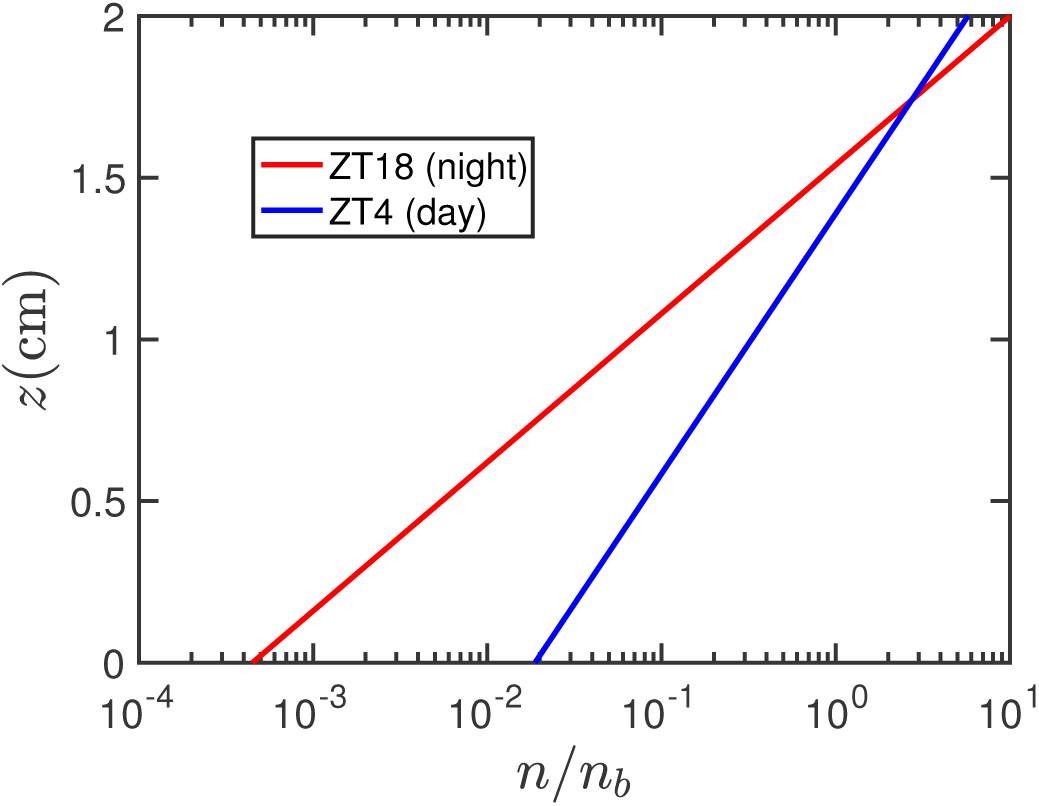
Predicted vertical distribution for *C. reinhardtii* in a capillary of height *h* = 2 cm, using motility parameters measured at ZT4 (light phase) and ZT18 (dark phase), as shown.

The observed diurnal variation in the motility statistics of *C. reinhardtii* has interesting consequences for its population-scale behaviour. Consider, for example, a *C. reinhardtii* suspension in a vertically oriented container that does not support large scale flows, e.g. a capillary. The transport in such a suspension is driven by fluxes due to gravitactic swimming, which drives cells to the top of the container, and anisotropic diffusion, which homogenises the distribution of cells (preferentially in the vertical direction). At long times these fluxes balance, and the steady state distribution of cells can be shown to be given by *n*(*z*) ∼ *e*^*z/l*^, where *z* is the vertical coordinate and *l* = *v*_s_ *D*_r_*f*(*λ*) is the characteristic length scale of the distribution, with *v*_s_ the swimming speed, *D*_r_ the rotational diffusivity and *f*(*λ*) a function of the bias parameter *λ*, derived from the biases in direction and diffusivity (see Materials and Methods for a detailed derivation). In previous studies, e.g. (14, 17, 25), these parameters have been considered fixed and independent of the growth illumination. Here, instead, we account for their diurnal variation and evaluate the length *l* with parameters measured in the light and dark phases of a diurnal cycle. For example, taking values from ZT4 and ZT18, we find *l*_*light*_ /*l*_*dark*_ = 0.6. That is to say, *C. reinhardtii* distributes vertically more broadly during the light than in the dark phase. This is clear from a plot of the steady state distributions plotted for ZT4 and ZT18, see Figure 4.

Diurnal variations also affect gyrotaxis, the orientational bias on swimming microalgae caused by a combination of gravitational and viscous torques, the latter due to shear in a flow. In related work, we use a combination of experiment and numerical modelling to study how the distribution of *C. reinhardtii* cells in a downwelling pipe flow changes as a function of shear and zeitgeber time in a diurnal cycle (Jin *et al.* in preparation). In future work, it would also be very interesting to connect diurnal variations in motility with the diel migrations of freshwater and marine phytoplankton. Future experiments could explore if the enhancement in swimming speed during the dark phase we have observed also occurs in other phytoplankton species, and test the consquences of this enhanced speed, which current models assume to be constant (39). Finally, building on the present work it would also be of great interest to quantitate diurnal variations in phototactic motility, revisiting the classic work by Bruce(8) using microfluidic experiments.

## AUTHOR CONTRIBUTIONS

O.A.C., D.J., and J.K conceived the experiments, D.J. and E.S. developed the data analysis, D.J. conducted the experiment, data analysis, and simulations. All authors reviewed the manuscript.

## ACKNOWLEDGMENTS

We thank Ms Dianyi Liu (Donald Danforth Plant Science Center, MO, USA) for suggestions on synchronisation to a light-dark cycle and discussion of the mitotic cell-cycle. We thank Prof. Stuart Dalziel (Department of Applied Mathematics and Theoretical Physics, University of Cambridge) for advice on cell tracking analysis. We thank Prof. Masahide Kikkawa and Dr Shohei Fujita (University of Tokyo) for kindly sharing the cell tracking data from their publication.

## REFERENCES

1. Lakin-Thomas, P. L., and S. Brody, 2004. Circadian Rhythms in Microorganisms: new complexities. Annual Review of Microbiology 58:489–519.

2. Mittag, M., S. Kiaulehn, and C. H. Johnson, 2005. The circadian clock in *Chlamydomonas reinhardtii*. What is it for? What is it similar to? Plant Physiology 137:399–409.

3. Strenkert, D., S. Schmollinger, S. D. Gallaher, P. A. Salomé, S. O. Purvine, C. D. Nicora, T. Mettler-Altmann, E. Soubeyrand, A. P. M. Weber, M. S. Lipton, G. J. Basset, and S. S. Merchant, 2019. Multiomics resolution of molecular events during a day in the life of *Chlamydomonas*. Proceedings of the National Academy of Sciences 116:2374–2383.

4. Cross, F. R., and J. G. Umen, 2015. The *Chlamydomonas* cell cycle. The Plant Journal 82:370–392.

5. Elgeti, J., R. G. Winkler, and G. Gompper, 2015. Physics of microswimmers—single particle motion and collective behavior: a review. Reports on Progress in Physics 78:056601.

6. Jeanneret, R., M. Contino, and M. Polin, 2016. A brief introduction to the model microswimmer *Chlamydomonas reinhardtii*. The European Physical Journal Special Topics 225:2141–2156.

7. Wingfield, J. L., and K.-F. Lechtreck, 2018. *Chlamydomonas* Basal Bodies as Flagella Organizing Centers. Cells 7.

8. Bruce, V. G., 1970. The Biological Clock in *Chlamydomonas reinhardi*. The Journal of Protozoology 17:328–334.

9. Byrne, T. E., M. R. Wells, and C. H. Johnson, 1992. Circadian Rhythms of Chemotaxis to Ammonium and of Methylammonium Uptake in *Chlamydomonas*. Plant Physiology 98:879–886.

10. Yoshimura, K., Y. Matsuo, and R. Kamiya, 2003. Gravitaxis in *Chlamydomonas* reinhardtii studied with novel mutants. Plant and Cell Physiology 44:1112–1118.

11. Garcia, M., S. Berti, P. Peyla, and S. RafaÃ, 2011. Random walk of a swimmer in a low-Reynolds-number medium. Physical Review E - Statistical, Nonlinear, and Soft Matter Physics 83:1–4.

12. Martinez, V. A., R. Besseling, O. A. Croze, J. Tailleur, M. Reufer, J. Schwarz-Linek, L. G. Wilson, M. A. Bees, and W. C. K. Poon, 2012. Differential dynamic microscopy: A high-throughput method for characterizing the motility of microorganisms. Biophysical Journal 103:1637–1647.

13. Hansen, T. J., M. Hondzo, M. T. Mashek, D. G. Mashek, and P. A. Lefebvre, 2013. Algal swimming velocities signal fatty acid accumulation. Biotechnology and Bioengineering 110:143–152.

14. Barry, M., 2014. Mechanisms of Reorientation in Phytoplankton: Fluid Shear, Surface Interactions, and Gravitaxis. Ph.D. thesis, Massachusetts Institute of Technology.

15. Fujita, S., T. Matsuo, M. Ishiura, and M. Kikkawa, 2014. High-throughput phenotyping of *Chlamydomonas* swimming mutants based on nanoscale video analysis. Biophysical Journal 107:336–345. http://dx.doi.org/10.1016/j.bpj.2014.05.033.

16. Barry, M. T., R. Rusconi, J. S. Guasto, and R. Stocker, 2015. Shear-induced orientational dynamics and spatial heterogeneity in suspensions of motile phytoplankton. Journal of The Royal Society Interface 12:20150791.

17. Croze, O. A., R. N. Bearon, and M. A. Bees, 2017. Gyrotactic swimmer dispersion in pipe flow: Testing the theory. Journal of Fluid Mechanics 816:481–506.

18. Sengupta, A., F. Carrara, and R. Stocker, 2017. Phytoplankton can actively diversify their migration strategy in response to turbulent cues. Nature 543:555–558.

19. Chen, X., Y. Wu, and L. Zeng, 2018. Migration of gyrotactic micro-organisms in water. Water 10:1455.

20. Harris, E. H., 1989. The Chlamydomonas Sourcebook: A Comprehensive Guide to Biology and Laboratory Use. Academic Press. https://www.amazon.com/{Chlamydomonas}-Sourcebook-Comprehensive-Biology-Laboratory/dp/012326880X?SubscriptionId=0JYN1NVW651KCA56C102&tag=techkie-20&linkCode=xm2&camp=2025&creative=165953&creativeASIN=012326880X.

21. Garz, A., M. Sandmann, M. Rading, S. Ramm, R. Menzel, and M. Steup, 2012. Cell-to-cell diversity in a synchronized *Chlamydomonas* culture as revealed by single-cell analyses. Biophysj 103:1078–1086. http://dx.doi.org/10.1016/j.bpj.2012.07.026.

22. Foster, K., and R. Smyth, 1980. Light Antennas in phototactic algae. Microbiological Reviews 44:572–630.

23. Crocker, J. C., and D. G. Grier, 1996. Methods of digital video microscopy for colloidal studies. Journal of Colloid and Interface Science 179:298–310.

24. Ruffer, U., and W. Nultsch, 1985. High-speed cinematographic analysis of the movement of *Chlamydomonas*. Cell motility 263:251–263.

25. Hill, N. A., and D. P. Hader, 1997. A biased random walk model for the trajectories of swimming micro-organisms. Journal of theoretical biology 186:503–526.

26. Pedley, T. J., Kessler, J. O., 1992. Hydrodynamic phenomena in suspensions of swimming microorganisms. Annual Review of Fluid Mechanics 24:313–358.

27. Pedley, T. J., and J. O. Kessler, 1990. A new continuum model for suspensions of gyrotactic micro-organisms. Journal of fluid mechanics 212:155–182.

28. Polin, M., I. Tuval, K. Drescher, J. P. Gollub, and R. E. Goldstein, 2009. *Chlamydomonas* swims with two “gears” in a eukaryotic version of run-and-tumble locomotion. Science 325:487.

29. Wan, K. Y., K. C. Leptos, and R. E. Goldstein, 2014. Lag, lock, sync, slip: the many ‘phases’ of coupled flagella. Journal of The Royal Society Interface 11:20131160.

30. Cavalier-Smith, T., 1974. Basal body and flagellar development during the vegetative cell cycle and the sexual cycle of *Chlamydomonas reinhardii*. Journal of Cell Science 16:529–56. http://www.ncbi.nlm.nih.gov/pubmed/4615103.

31. Zones, J. M., I. K. Blaby, S. S. Merchant, and J. G. Umen, 2015. High-resolution profiling of a synchronized diurnal transcriptome from *Chlamydomonas reinhardtii* reveals continuous cell and metabolic differentiation. The Plant Cell 27:2743–2769. http://www.plantcell.org/content/early/2015/10/02/tpc.15.00498.abstract?papetoc.

32. Guasto, J. S., K. A. Johnson, and J. P. Gollub, 2010. Oscillatory flows induced by microorganisms swimming in two dimensions. Physical Review Letters 105:18–21.

33. Roberts, A. M., 2006. Mechanisms of gravitaxis in *Chlamydomonas*. Biological Bulletin 210:78–80.

34. Chen, D. T. N., M. Heymann, S. Fraden, D. Nicastro, and Z. Dogic, 2015. ATP consumption of eukaryotic flagella measured at a single-cell level. Biophysical Journal 109:2562–2573. http://dx.doi.org/10.1016/j.bpj.2015.11.003.

35. Raven, J. A., 1982. The energetics of freshwater algae: energy requirements for biosynthesis and volume regulation. New Phytologist 92:1–20.

36. Martínez-García, E., P. I. Nikel, M. Chavarría, and V. de Lorenzo, 2014. The metabolic cost of flagellar motion in Pseudomonas putida KT2440. Environmental Microbiology 16:291–303.

37. Lindemann, C. B., 2003. Structural-functional relationships of the dynein, spokes, and central-pair projections predicted from an analysis of the forces acting within a flagellum. Biophysical Journal 84:4115–4126.

38. Johnson, X., and J. Alric, 2013. Central carbon metabolism and electron transport in *Chlamydomonas* reinhardtii: Metabolic constraints for carbon partitioning between oil and starch. Eukaryotic Cell 12:776–793.

39. Richards, S. A., H. P. Possingham, and J. Noye, 1996. Diel vertical migration: modelling light-mediated mechanisms. Journal of Plankton Research 18:2199–2222.

